# Deciphering the roadmap of *in vivo* reprogramming towards pluripotency

**DOI:** 10.1101/2022.04.19.488763

**Authors:** Dafni Chondronasiou, Jaime Martínez de Villareal, Elena Melendez, Cian J. Lynch, Marta Kovatcheva, Mònica Aguilera, Neus Prats, Francisco X. Real, Manuel Serrano

**Affiliations:** Institute for Research in Biomedicine (IRB Barcelona), Barcelona Institute of Science and Technology (BIST), Barcelona, Spain; Epithelial Carcinogenesis Group, Molecular Oncology Programme, Spanish National Cancer Research Centre (CNIO), Madrid, Spain; CIBERONC, Madrid, Spain; Departament de Medicina i Ciències de la Vida, Universitat Pompeu Fabra, Barcelona, Spain; Catalan Institution for Research and Advanced Studies (ICREA), Barcelona 08010, Spain

**Keywords:** Reprogramming, OSKM, pluripotency, dedifferentiation, pancreas

## Abstract

Differentiated cells can be converted to pluripotent stem cells (iPSCs) upon ectopic expression of transcription factors OCT4, SOX2, KLF4 and MYC (OSKM) in a process known as reprogramming. Great efforts have been made to dissect intermediate states of *in vitro* reprogramming and how they are affected by culture conditions, while the roadmap of *in vivo* reprogramming remains unexplored. Here, we use single cell RNA sequencing to capture cells undergoing reprogramming in the adult pancreas. We identify markers along the trajectory from acinar identity to pluripotency, which allow *in situ* visualization of the intermediate states of reprogramming. Importantly, different tissues expressing OSKM, such as pancreas, stomach and colon, share markers of intermediate reprogramming, suggesting a conserved *in vivo* reprogramming path. Our *in vivo* roadmap defines landmarks along *in vivo* reprogramming that could be useful for applications in tissue regeneration and cellular rejuvenation based on intermediate reprogramming states.

## Introduction

The ability to manipulate cell fate *in vitro* has revolutionized regenerative medicine. The most striking breakthrough in the field occurred when Yamanaka first illustrated the ability of adult differentiated cells to give rise to pluripotent cells (induced pluripotent stem cells or iPSCs) upon the simultaneous expression of four transcriptional factors, OCT4, SOX2, KLF4 and MYC (OSKM), also known as “Yamanaka factors” (Takahashi and Yamanaka, 2006). Through this process of reprogramming, a fraction of adult somatic cells are able to shut down the transcriptional programs linked to their cell identity and progressively activate the transcriptional network of pluripotency (Deng et al., 2021; Takahashi and Yamanaka, 2006). In mice, OSKM expression recapitulates the events of cellular dedifferentiation in multiple tissues and leads to the emergence of pluripotent cells (Abad et al., 2013; Mosteiro et al., 2016; Ohnishi et al., 2014). The ultimate manifestation of complete reprogramming *in vivo* is the formation of teratomas, a tumor originated from iPSCs differentiating into all three germ layers (Abad et al., 2013; Ohnishi et al., 2014).

Cellular reprogramming is an inefficient process due, at least in part, to the existence of multiple cell-autonomous barriers, such as tumour suppressors, chromatin regulators, transcription factors, signalling pathways and micro RNAs (Arabacı et al., 2021; Haridhasapavalan et al., 2020). Various studies have tried to untangle the complex cascade of molecular and epigenetic events occurring during *in vitro* reprogramming as well as to define intermediate states (Brambrink et al., 2008; Chronis et al., 2017; O’Malley et al., 2013; Polo et al., 2012; Stadtfeld et al., 2008; Zviran et al., 2019). The vast majority of cells do not successfully complete reprogramming and have varying cell fates: some undergo apoptosis, mainly triggered by MYC induction (Kim et al., 2018), while others senesce as a result of the activation of two tumor suppressor pathways, p53 and INK4A/ARF (Banito et al., 2009; Hong et al., 2009; Kawamura et al., 2009; Li et al., 2009; Marión et al., 2009; Utikal et al., 2009). On the other hand, senescent cells contribute in a paracrine manner to *in vitro* reprogramming through IL6 secretion (Brady et al., 2013; Chiche et al., 2017; Mosteiro et al., 2016). Another alternative OSKM-induced cellular outcome *in vitro* is a pluripotent state epigenetically and transcriptionally distinct from iPSCs designated as “F-class” cells because of their fuzzy appearance (Benevento et al., 2014; Clancy et al., 2014; Hussein et al., 2014; Tonge et al., 2014). Cells with placenta, neuronal or epidermal-like identities have also been described as reprogramming intermediates *in vitro* (Kurian et al., 2013; O’Malley et al., 2013; Schiebinger et al., 2017). Additionally, trophectoderm (TSCs) and extraembryonic endoderm (XEN) were also reported as alternative trajectories of OSKM reprogramming, illustrating the heterogeneity of this process (Deng et al., 2021; Liu et al., 2020; Parenti et al., 2016; Schiebinger et al., 2019; Xing et al., 2020). Finally, intermediate reprogrammed fibroblasts are able to generate mesodermal progenitors upon culturing with the adequate medium, which can subsequently differentiate into endothelium and smooth muscle (Kurian et al., 2013).

However, very little is known about the molecular roadmap of *in vivo* reprogramming. Here, we used single cell transcriptomics to further decipher the process of reprogramming *in vivo*. Focusing on pancreas, as the tissue with the highest reprogramming efficiency in our mouse model (Abad et al., 2013; Mosteiro et al., 2016), we identify markers of intermediate reprogramming and map those markers into the reprogrammed tissue using immunohistochemistry and RNA fluorescence *in situ* hybridization. Moreover, we demonstrate that many of those markers are shared among other tissues reprogrammed *in vivo* and mouse embryonic fibroblasts (MEFs) reprogrammed *in vitro*. To our knowledge, this is the first time that an OSKM-driven roadmap has been generated *in vivo* in the context of an adult organism.

## Results

### scRNAseq captures partially reprogrammed populations in the pancreas

To capture the cellular heterogeneity generated during partial reprogramming in the pancreas, we performed single cell RNA sequencing (scRNAseq). Three reprogrammable (OSKM) and two control (WT) mice were treated with doxycycline (1 mg/ml) for 7 days (**Figure 1A**). Upon examination of the pancreas, we could appreciate that the extent of reprogramming was higher in two individual OSKM mice, mice OSKM 2 and 3, and modest in mouse OSKM 1. For comparison, we treated another group of mice with cerulein (CER) for 2 consecutive days (**Figure 1A**). Cerulein produces severe acinar damage leading to acinar cell plasticity and acinar-to-ductal metaplasia (ADM). At the end of the treatment, each pancreas was dissociated into single-cell suspension, sorted for live (DAPI negative) cells, and sequenced using the 10x Genomics system. From each individual pancreas, we sequenced between 6,657 to 14,240 cells, amounting to a total of 52,717 cells, which we analyzed with the R package Seurat. All samples were merged without integration (**Figure S1A,D, S2A**), clustered, and manually annotated based on the expression of defined markers (see Experimental Procedures) (**Figure S1B,C,E,F, S2B,C, Table S1**). We observed that in all three conditions, even in the context of reprogramming, all main pancreatic cell types could still be identified, which is consistent with the fact that reprogramming and cerulein-induced ADM are focal, not affecting the entire pancreas. However, cell type composition was dramatically altered after OSKM activation or cerulein damage, as was clearly depicted in the Uniform Manifold Approximation and Projection (UMAP) by the abundant infiltration of inflammatory cells, activated fibroblasts likely corresponding to pancreatic stellate cells, and the under-representation of epithelial cells (dropping from 45.5% in WT pancreas to 10.5% and to 15.3% in OSKM and CER pancreas, respectively) (**Figure S1, S2**). Of note, an analysis of the inflammatory cells present during reprogramming can be found elsewhere (Melendez et al., 2022). Here, we have focused on the epithelial compartment of the pancreas, as it is the most extensively altered histologically (**Figure 1A**).

**Figure 1.**
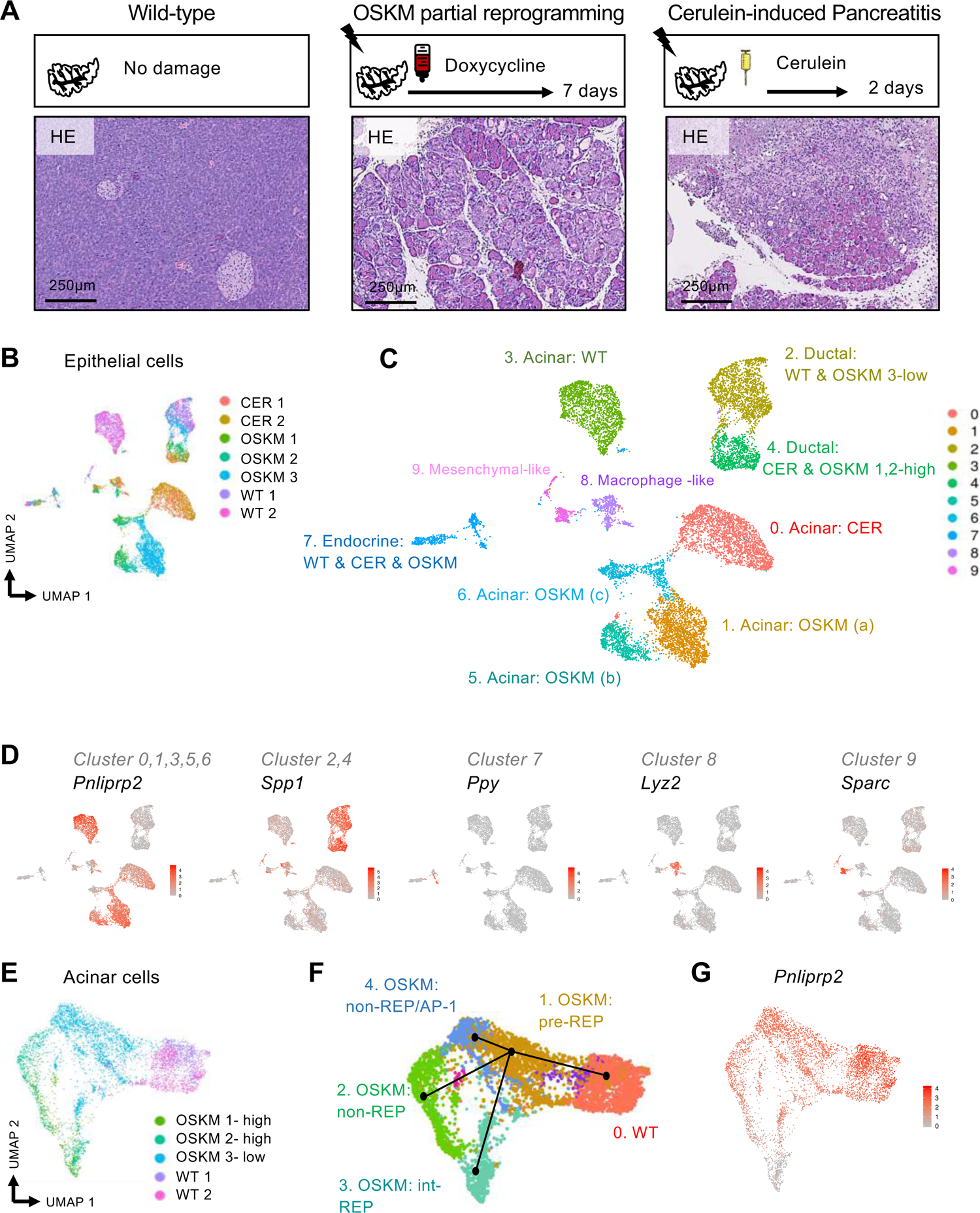
Single cell RNA sequencing reveals the susceptibility of pancreatic acinar cells to *in vivo* OSKM reprogramming. (A) Representative images of Hematoxylin/Eosin (HE) staining of pancreas from wild-type (WT), cerulein (CER)-treated (total of 14 i.p. injections in two consecutive days) for inducing acute pancreatitis, and OSKM-induced mice (1mg/ml doxycycline in the drinking water for 7 days) for inducing partial reprogramming. (B) All epithelial cells (acinar, ductal, endocrine) coming from all samples (two WT, two CER and three OSKM mice) were selected, merged and reanalyzed as represented in this UMAP (Uniform Manifold Approximation and Projection). The contribution of each sample is depicted in this UMAP by a different color. (C) Merged cells were clustered using a cluster resolution parameter of 0.2 and each cluster was annotated according to the expression of characteristic markers. (D) Established marker genes identify acinar cells (*Pnliprp2*), ductal cells (*Spp1*), endocrine cells (*Ppy*), macrophage-like (*Sparc*) and mesenchymal-like (*Sparc*) cells. (E) All acinar-related clusters coming from WT and OSKM-induced mice (*clusters 1,3,5,6*) were selected, merged and reanalyzed as represented in this UMAP. The contribution of each sample is depicted in this UMAP by a different color. (F) Trajectory analysis of the acinar cells coming from WT and OSKM-induced mice using the Slingshot tool (Street et al., 2018). (G) UMAP visualization of the acinar marker *Pnliprp2* in acinar cells corresponding to WT and OSKM pancreata.

To capture the differences among WT, CER-treated and OSKM-induced pancreas at epithelial level, we selected acinar, ductal and endocrine cells coming from all samples, and we reanalyzed them (**Figure 1B**). Acinar gene expression was detected in 5 independent clusters (WT: cluster 3; CER: cluster 0, OSKM: clusters 1, 5 and 6) (**Figure 1C,D, Table S2**). On the other hand, ductal cells coming from pancreas OSKM 2 and 3 clustered together with CER ductal cells (cluster 4), but not with ductal cells from control mice, suggesting that damage induced by cerulein or OSKM affects ductal cells in similar ways. Finally, endocrine cells clustered together (cluster 7) regardless of their origin (**Figure 1C,D, Table S2**). Overall, the epithelial analysis of the scRNAseq data suggest that acinar cells are the most affected in the pancreas after OSKM activation in our mouse model. Moreover, OSKM expression produces distinct transcriptional changes in the acinar compartment compared to cerulein-induced acute pancreatitis.

To gain further insight into the acinar-associated transcriptional states generated by OSKM activation, we extracted only acinar cells coming from OSKM and WT pancreas (**Figure 1E**). After reclustering, we identified 5 acinar clusters, one for WT and four for OSKM, and we performed cell lineage and pseudotime trajectory inference analysis using Slingshot (Street et al., 2018), pre-setting WT acinar cells as starting point (**Table S3**). In this way, we identified three main trajectories (or *pseudotimes*) emerging from a common OSKM cluster (**Figure 1F, S2D**). We will refer to this precursor cluster as pre-reprogrammed or pre-REP (**Figure 1F**). This pre-REP cluster is the hub for the emergence of three different trajectories. Two of them retain acinar identity and therefore we considered them as “non-reprogrammed” fates or non-REP (**Figure 1F**). We noted that one of these clusters was characterized by an abundant presence of the transcription factors AP-1, so we refer to this cluster as non-REP/AP-1 (**Figure 1F**). Interestingly, a third trajectory emerging from pre-REP gives rise to a cluster of cells that have largely lost their acinar identity, as judged by the strong reduction of acinar markers (**Table S3**), such as *Pnliprp2* (**Figure 1G**), which we refer to it as intermediate-reprogrammed cells or int-REP (**Figure 1F**).

### *In vivo* identification of “pre-reprogramming” and “non-reprogramming” OSKM fates

Upon examination of the pre-REP and non-REP clusters (**Tables S2, S3**), we focused on *Reg3a* and related family members because they were highly expressed in both clusters (**Figure 2A, S3A**) and, also, to take advantage of the availability of antibodies for immunohistochemistry. The “regenerating” (Reg) family of C-type lectins is known to be upregulated in pancreatitis (Chen et al., 2019) and, indeed, we observed upregulation of several *Reg3*-family members in cerulein-treated pancreas, but the level of upregulation was much lower compared to OSKM-expressing pancreas (**Figure 2B, S3A**). We visualized the presence of REG3A/G-positive cells in pancreas undergoing reprogramming by immunohistochemistry using an antibody that recognizes these two members of the REG3 family (**Figure 2C**). Interestingly, REG3A/G cells were rare in the dysplastic areas with abnormal histology, but abundant in the regions of the exocrine pancreas with apparently normal histology (**Figure 2C**). This suggests that the pre-REP and non-REP cells retain a normal histology while undergoing transcriptional changes.

**Figure 2.**
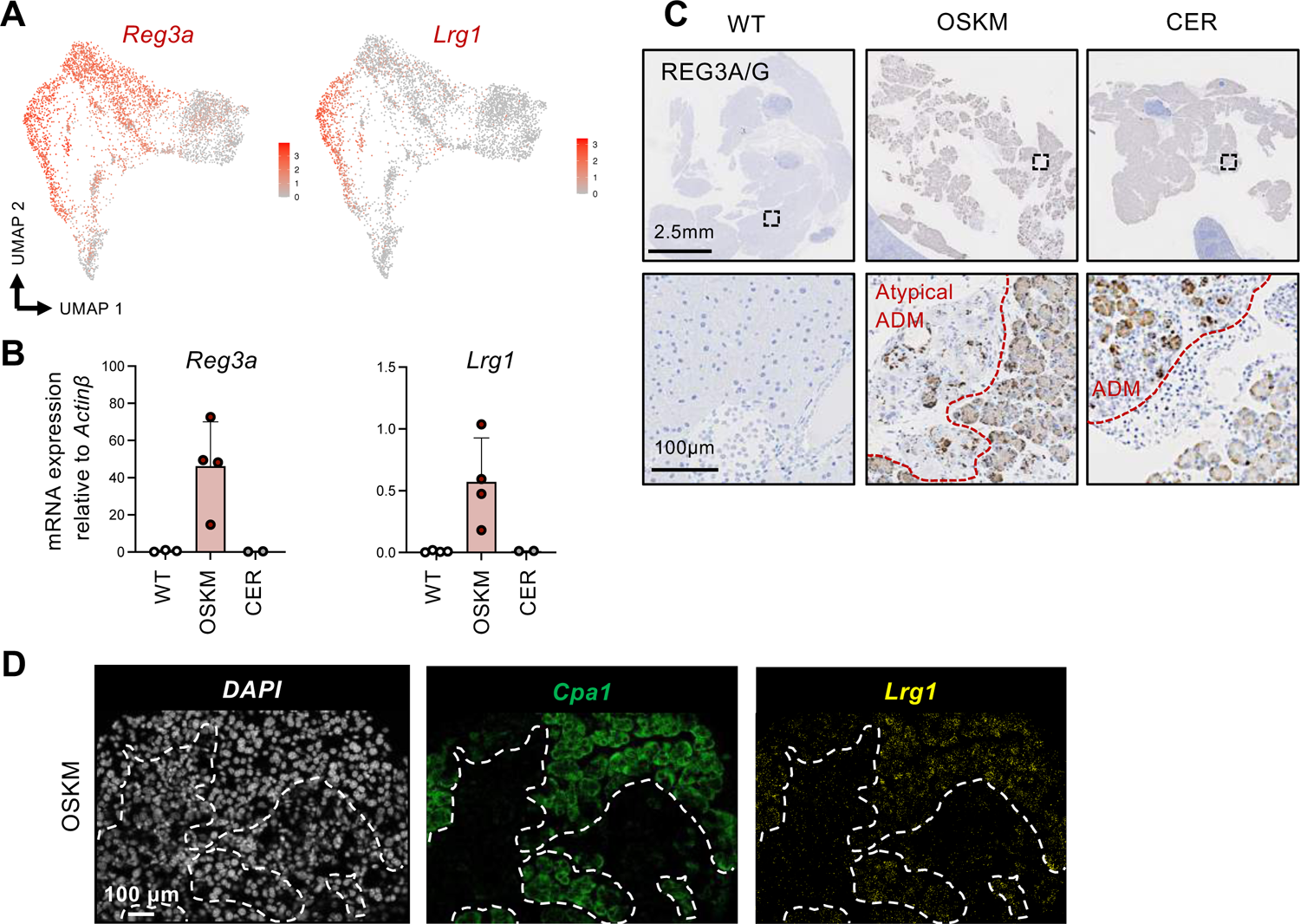
*In vivo* identification of “pre-reprogramming” and “non-reprogramming” OSKM fates. (A) UMAP representation of *Reg3a* and *Lrg1* expression in acinar cells coming from WT and OSKM-induced pancreata. (B) mRNA expression of *Reg3a* and *Lrg1* in bulk pancreas comparing all three conditions. (C) Immunohistochemistry of REG3A/G in the pancreas of WT, CER-treated and OSKM-induced mice. (D) RNA fluorescence *in situ* hybridization (RNA-FISH) of *Cpa1* (acinar marker) and *Lrg1* in reprogrammed pancreas. Scale bar, 100 μm.

The pre-REP and non-REP clusters shared most of their markers, with the exception of a few ones that were exclusive of the non-REP cluster, being even absent from the related non-REP/AP-1 cluster. This is the case of *Lrg1* (leucine-rich alpha-2-glycoprotein 1) and *Lcn2* (lipocalin 2), which were highly upregulated in the non-REP cluster (**Figure 2A, S3B, Tables S2, S3**). Interestingly, qRT-PCR of bulk pancreas RNA revealed that this gene was dramatically upregulated in pancreas upon OSKM expression but not upon cerulein treatment (**Figure 2B**). RNA fluorescence *in situ* hybridization (RNA-FISH) indicated that *Lrg1* expression was restricted to pancreatic areas that maintained their acinar identity (*Cpa1*-positive) in the context of OSKM-expressing pancreas (**Figure 2D**). Another gene exclusively expressed in the non-REP cluster is *Lcn2* (**Figure S3B**).

Finally, we wondered if non-REP cells were actively expressing the transgenic OSKM cassette. By RNA-FISH, we observed that the expression of transgenic OSKM after 7 days of doxycycline was largely restricted to the pancreas regions that had lost acinar identity (*Cpa1* negative) (**Figure S3C**). These observations suggest that the markers of non-REP cells are not a cell-autonomous consequence of OSKM expression, but probably reflect the altered microenvironment caused by nearby OSKM-expressing cells.

### *In vivo* identification of “intermediate-reprogramming”

We then focused on the int-REP cluster. This cluster consists of a heterogeneous population of cells expressing genes that are not normally expressed in the pancreas. This cluster also contains a few cells expressing the pluripotency marker *Oct4* (**Figure S4**), reinforcing the idea that this cluster represents cells undergoing reprogramming. Examining the genes upregulated in the int-REP cluster (from the acinar UMAP in Fig.1F and from the epithelial UMAP cluster 6 in Fig. 1C), we selected 20 genes whose expression pattern was restricted to the int-REP cluster (**Figure S4, Table S2,3**). We evaluated their expression by RT-qPCR in bulk pancreas and we validated that almost all of them were selectively induced upon OSKM reprogramming, but not in cerulein-treated pancreas (**Figure 3A**). Based on their expression pattern (**Figure S4**), we could divide them into three categories that reflect their sub-localization along the reprogramming trajectory: early, advanced, or in both early and advanced sub-localizations (**Figure 3A, S4**). We noticed that *Muc5ac*, which is normally expressed in the gastric and respiratory tract, was early induced upon OSKM activation (**Figure 3B**). RNA-FISH illustrates the expression of *Muc5ac* both in a subset of cells that retain acinar identity (*Cpa1* positive) and in a subset of cells that have lost acinar identity (*Cpa1* negative) (**Figure 3C**).

**Figure 3.**
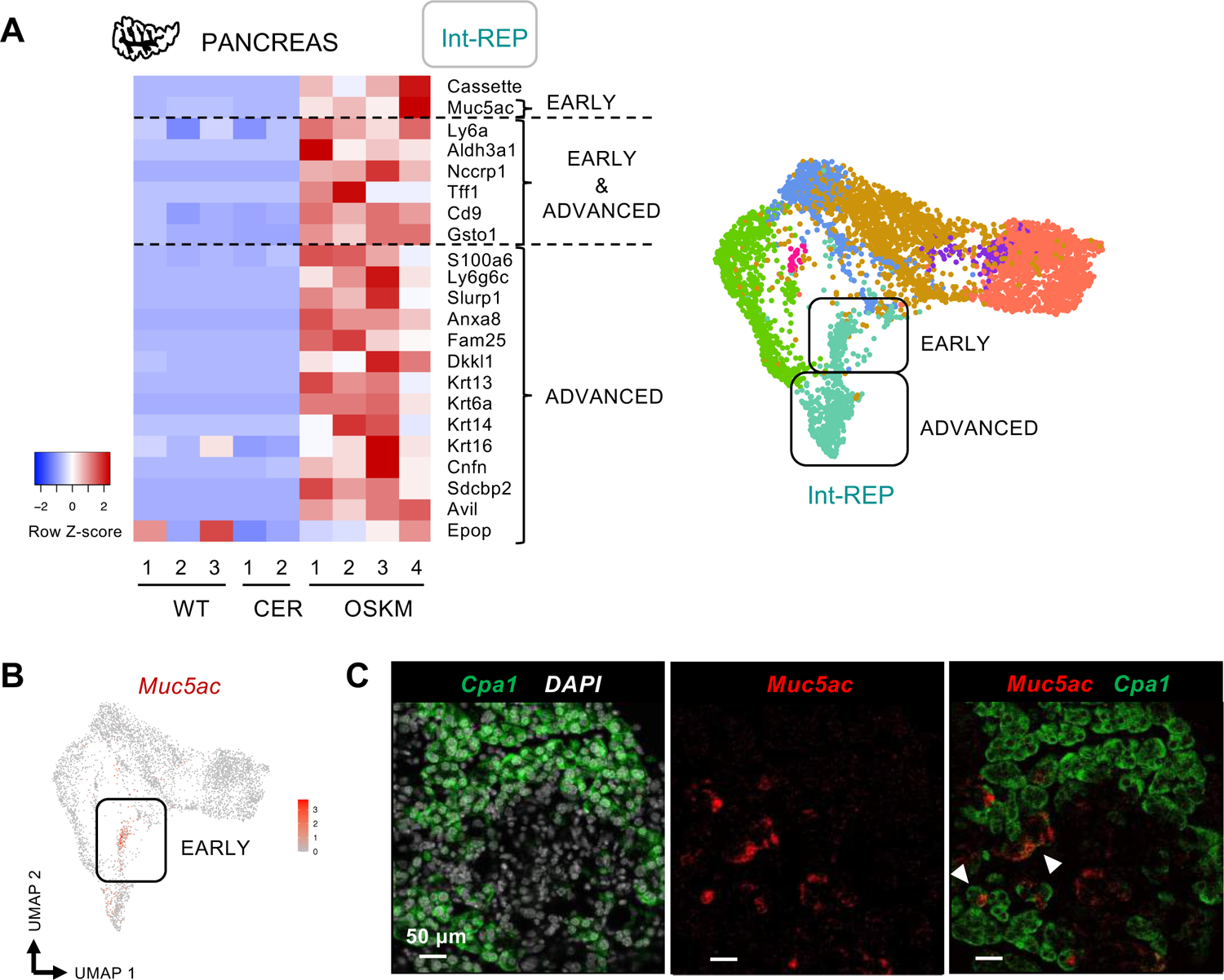
*In vivo* identification of “intermediate-reprogramming”. (A) Heatmap representation of mRNA expression measured by qRT-PCR of selected genes characterizing cells undergoing partial reprogramming in pancreas from WT (n=3), CER-treated (n=2) and OSKM-induced (n=4) mice. Blue represents low expression and red high expression of those genes. Depending on their expression pattern, genes have been divided into three groups: early, advanced or early & advanced. (B) UMAP representation of the expression of the “early” marker *Muc5ac* in acinar cells coming from WT and OSKM-induced pancreata. (C) Images of RNA-fluorescence *in situ* hybridization (RNA-FISH) of partially reprogrammed pancreas using probes against *Cpa1* and *Muc5ac* together with DAPI. Scale bars, 50 μm.

Among the “early and advanced” markers of int-REP cells, we noticed *Ly6a* (*Sca-1*) (**Figure 4A**), a member of the LY6 family associated with stem/progenitor cells in various tissues including pancreas (Holmes and Stanford, 2007; Leinenkugel et al., 2022) and previously reported to be upregulated during *in vitro* reprogramming of mouse embryonic fibroblasts (MEFs) as well (Schwarz et al., 2018). Remarkably, LY6A immunohistochemistry clearly defined the dysplastic regions of reprogrammed pancreas and the amount of LY6A+ cells was positively correlated with the extension of tissue dysplasia (**Figure 4B-D**). Immunohistochemistry did not detect LY6A+ cells in normal pancreas or in cerulein-treated pancreas (**Figure 4B**). Similarly, *Aldh3a1* was also upregulated in the entire int-REP cluster (early and advanced) (**Figure 4A**), and RNA-FISH revealed that it was largely expressed in cells that had lost acinar identity (*Cpa1* negative cells), although some cells co-expressed *Aldh3a1* and *Cpa1* (**Figure 4E**). *Aldh3a1* is a member of aldehyde dehydrogenase (ALDH) superfamily that is expressed in cornea; it is upregulated in response to oxidative stress, acting as an antioxidant (Vasiliou and Nebert, 2005) and participating in cell cycle regulation (Estey et al., 2007). Additionally, *Cd9* was another characteristic marker of this cluster (**Figure S5**), which has been previously identified as a marker of pancreatic cancer stem cells (Wang et al., 2019). Of note, we observed CD9 to be localized in the nucleus of OSKM dysplastic pancreatic cells (**Figure S5**). In this regard, a nuclear CD9 pool has previously been reported in the context of human breast carcinoma, where it participates in mitotic processes (Rappa et al., 2014).

**Figure 4.**
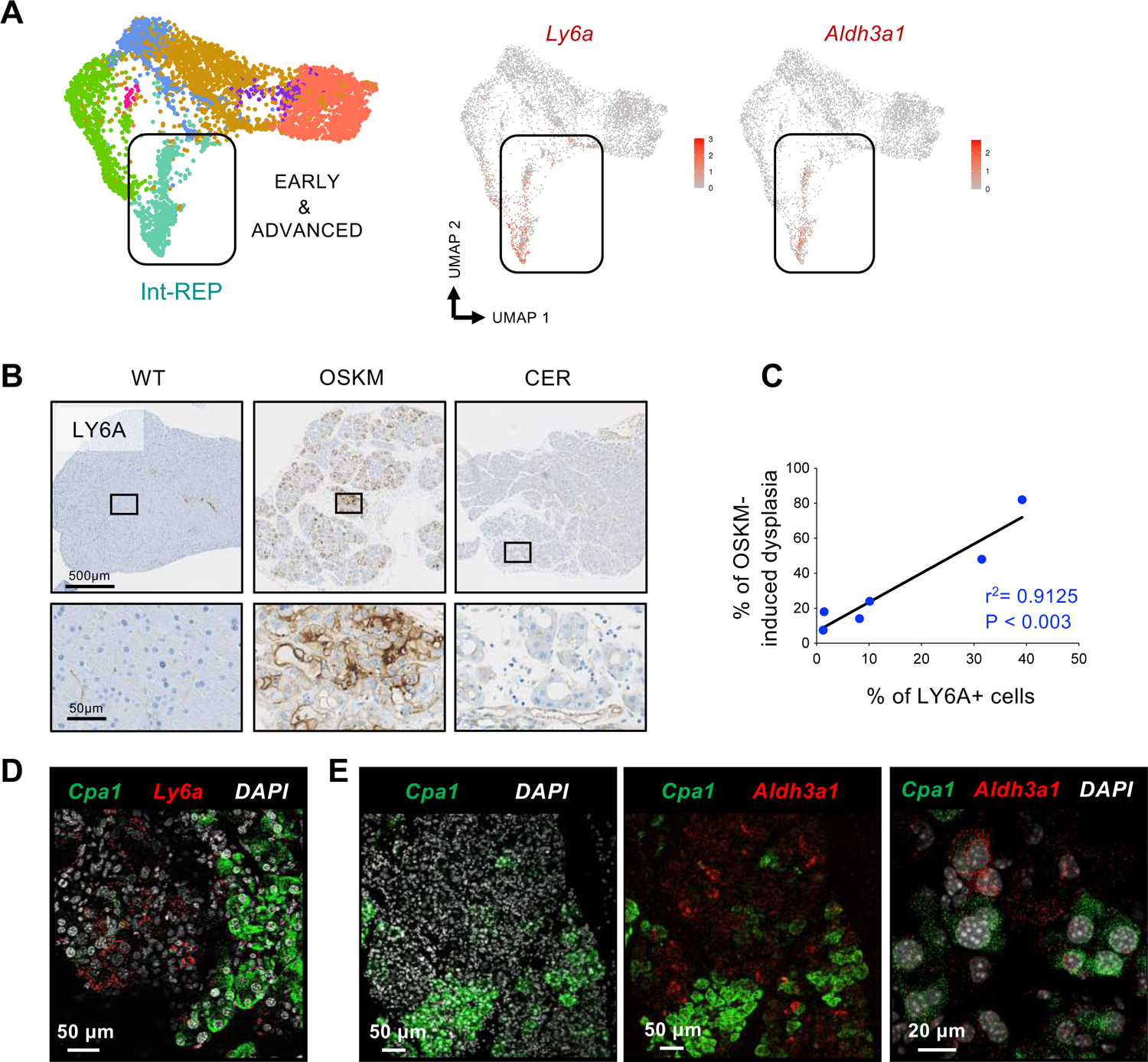
*In vivo* identification of early and advanced markers of “intermediate-reprogramming”. (A) UMAP representation of *Ly6a* and *Aldh3a1* expression in acinar cells coming from WT and OSKM-induced pancreata. (B) Immunohistochemistry of LY6A in paraffin-embedded sections of pancreas from WT, CER-treated and OSKM-induced mice. Scale bars, 500μm and 50μm. (C) The extent of OSKM-induced dysplasia is highly correlated with the presence of LY6A+ cells in the reprogrammed pancreas. (D) Image of RNA-fluorescence *in situ* hybridization (RNA-FISH) of partially reprogrammed pancreas using probes against *Cpa1* and *Ly6a* together with DAPI. (E) Images of RNA-FISH of partially reprogrammed pancreas using probes against *Cpa1* and *Aldh3a1* together with DAPI. Scale bars, 50μm and 20μm as indicated on each image separately.

Finally, we analyzed advanced int-REP genes. We noticed that a high number of keratins were upregulated in this group of genes, including those characteristics of stratified epithelia, such as *Krt14* (**Figure 5A**). Regions of dysplastic pancreas were clearly stained with KRT14, whereas WT and CER pancreas were completely negative (**Figure 5B**). *Krt14* is a keratin that it is not normally expressed in pancreas except in the context of rare squamous pancreatic cancers (Bailey et al., 2016; Real et al., 1993). Notably, the extent of OSKM-induced dysplasia was positively correlated with KRT14 expression in the pancreas (**Figure 5C**). Another gene in the advanced int-REP category is *Ly6g6c* which belongs to the LY6 family, although located in a different gene cluster than *Ly6a* (Upadhyay, 2019). Using RNA-FISH, we could map the *Ly6g6c* expressing cells into the de-differentiated areas where acinar cells had completely lost their identity (*Cpa1* negative) (**Figure 5D**). The partial co-localization of *Krt14* with *Ly6g6c* suggests that they are part of the same transcriptional trajectory, while their partial co-staining reflects the dynamic nature of the reprogramming process (**Figure 5D**). Other interesting genes in the advanced-intermediate reprogramming were *Slurp1* and *Sdcbp*. SLURP1 is another member of the LY6 family located in the same cluster as LY6A. SLURP1 is characterized by being secreted and acts as an agonist of cholinergic receptors and stabilizes epithelial cell-cell junctions (Campbell et al., 2019; Vasilyeva et al., 2017).

**Figure 5.**
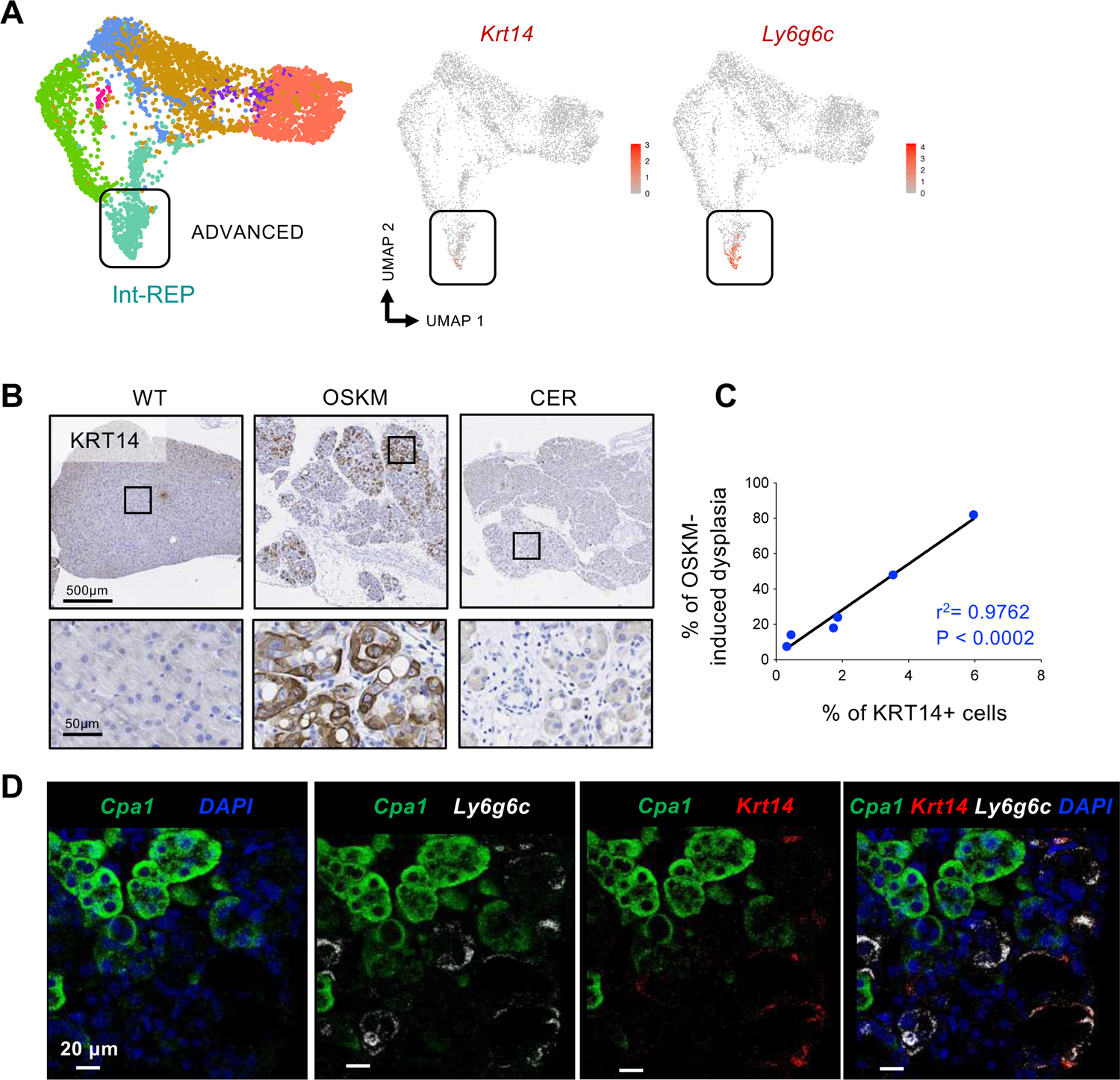
*In vivo* identification of advanced markers of “intermediate-reprogramming”. (A) UMAP representation of *Krt14* and *Ly6g6c* expression in acinar cells coming from WT and OSKM-induced pancreata. (B) Immunohistochemistry of KRT14 in paraffin-embedded sections of pancreas from WT, CER-treated and OSKM-induced mice. Scale bars, 500μm and 50μm. (C) The extent of OSKM-induced dysplasia is highly correlated with the presence of KRT14+ cells in the reprogrammed pancreas. (D) Images of RNA-fluorescence *in situ* hybridization (RNA-FISH) of partially reprogrammed pancreas using probes against *Cpa1*, *Ly6g6c* and *Krt14* together with DAPI. Scale bars, 20μm.

### A roadmap of *in vivo* reprogramming in the pancreas

To further refine and support the existence of early and advanced states of intermediate reprogramming, we performed a number of additional combined RNA-FISH staining. In particular, we observed that *Muc5ac* and *Ly6g6c*, which were classified as markers of early- and advanced-intermediate reprogramming, respectively, did not co-localize (**Figure 6A**). Regarding *Ly6g6c*, we have shown extensive co-localization with *Krt14* (**Figure 5D**), which is consistent with the concept that both genes are markers of advanced-intermediate reprogramming. In contrast *Krt14*, co-localized with a subset of *Aldh3a1*, a marker that spans both early- and advanced-intermediate reprogramming (**Figure 6B**). Collectively, our data support a first roadmap for *in vivo* reprogramming in the pancreas that is supported by single-cell RNAseq data, qRT-PCR in bulk tissues, and RNA-FISH and immunohistochemistry in histological sections (**Figure 6C**).

**Figure 6.**
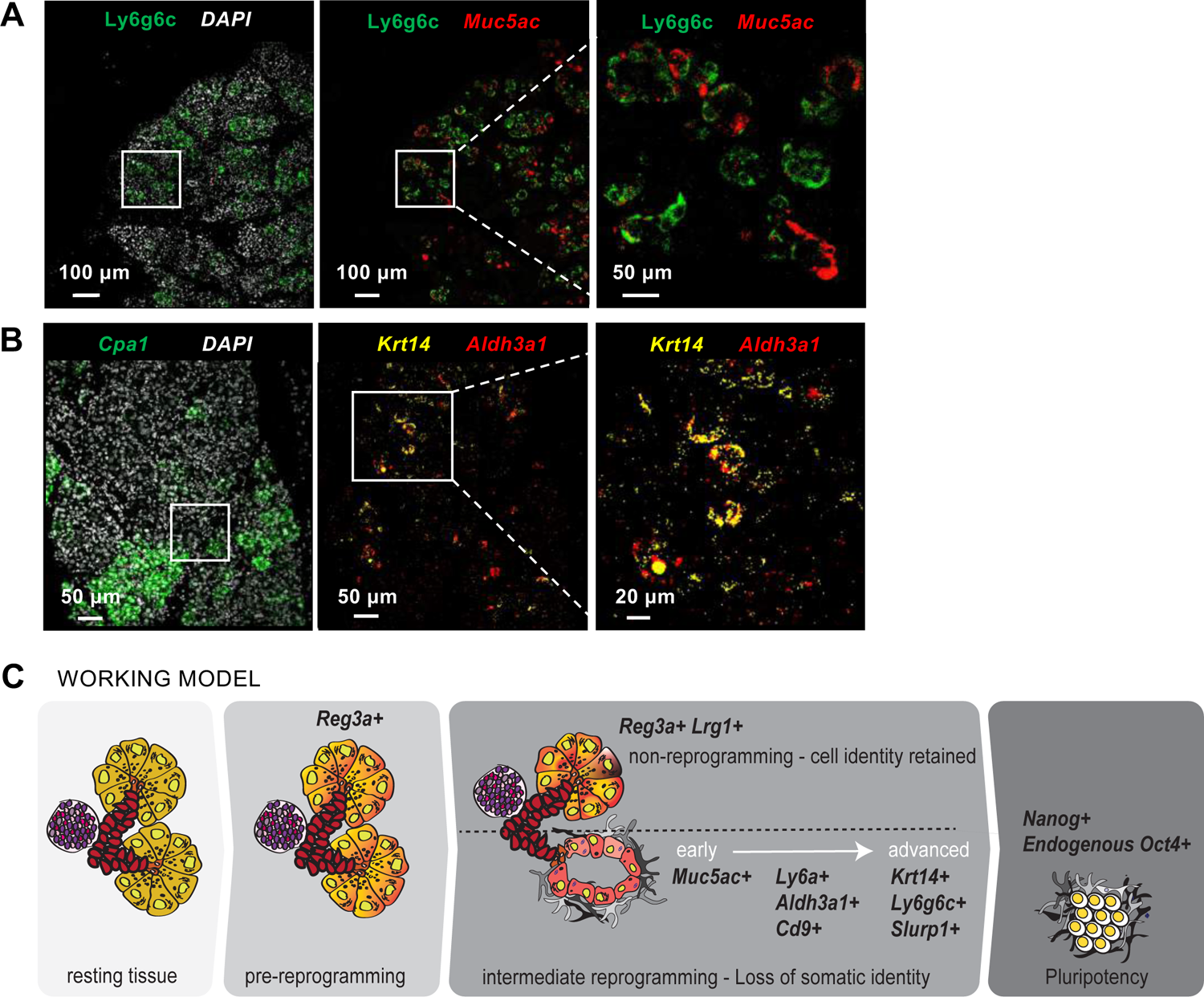
A roadmap of *in vivo* reprogramming in the pancreas. (A) RNA-FISH visualizes the expression of multiple intermediate-reprogramming markers in OSKM-induced pancreas using probes: (A) against *Ly6g6c* and *Muc5ac* together with DAPI, (B) against *Cpa1*, *Aldh3a1* and *Krt14* together with DAPI. Section bars, 100μm and 50μm as indicated on each image separately. (C) Our working model illustrates the molecular roadmap induced by OSKM activation in pancreas. Briefly, upon OSKM activation *in vivo*, whole acini react by upregulating *Reg3a* (“pre-reprogramming” state). Subsequently, while a fraction of acinar cells are refractory to reprogramming and retain their identity (*Reg3a*+ *Lrg1*+: “non-reprogramming” state), another fraction loses its acinar identity and acquires distinct gene expression profiles along the process of dedifferentiation (“intermediate-reprogramming”). Ultimately, few cells successfully make it to the pluripotent state, marked by the expression of *Nanog* and endogenous *Oct4*.

### Markers of “intermediate reprogramming” are shared among several tissues

Having identified markers of intermediate OSKM reprogramming in the exocrine pancreas, we wondered if these markers would also be upregulated during the reprogramming of other cell types. We first examined their expression by qRT-PCR in primary mouse embryo fibroblasts (MEFs) undergoing *in vitro* reprogramming. Remarkably, all of the tested pancreatic int-REP markers were upregulated during *in vitro* reprogramming of MEFs and the majority were absent in the final state of iPS (**Figure S6A**). Given the conservation of markers between these different reprogramming models, we evaluated their expression in the stomach and colon of OSKM-induced mice (**Figure 7A**). Interestingly, several genes of this signature were also upregulated by qRT-PCR in reprogrammed stomach and colon compared to WT samples (**Figure 7A**). By immunohistochemistry, dysplastic epithelial cells in reprogramming glandular stomach and colon were clearly stained by LY6A and KRT14 after 7 days of treatment with doxycycline, while normal WT tissues did not express these markers (**Figure 7B**). Of note, it has been previously reported that in the context of ulcerative colitis, the regenerating intestinal epithelium acquires a fetal-like expression profile marked by LY6A expression (Yui et al., 2018). Another interesting observation is that the non-dysplastic epithelial regions of the reprogramming intestine were positive for REG3A/G immunohistochemistry, similar to the non-reprogramming areas of the pancreas upon OSKM expression (**Figure S6B**). In case of the glandular stomach, expression of OSKM also clearly induced REG3A/G+ cells, although their number was clearly lower than in pancreas or intestine. Therefore, non-REP and int-REP markers originally identified in pancreas behave in a similar manner in stomach and colon.

**Figure 7.**
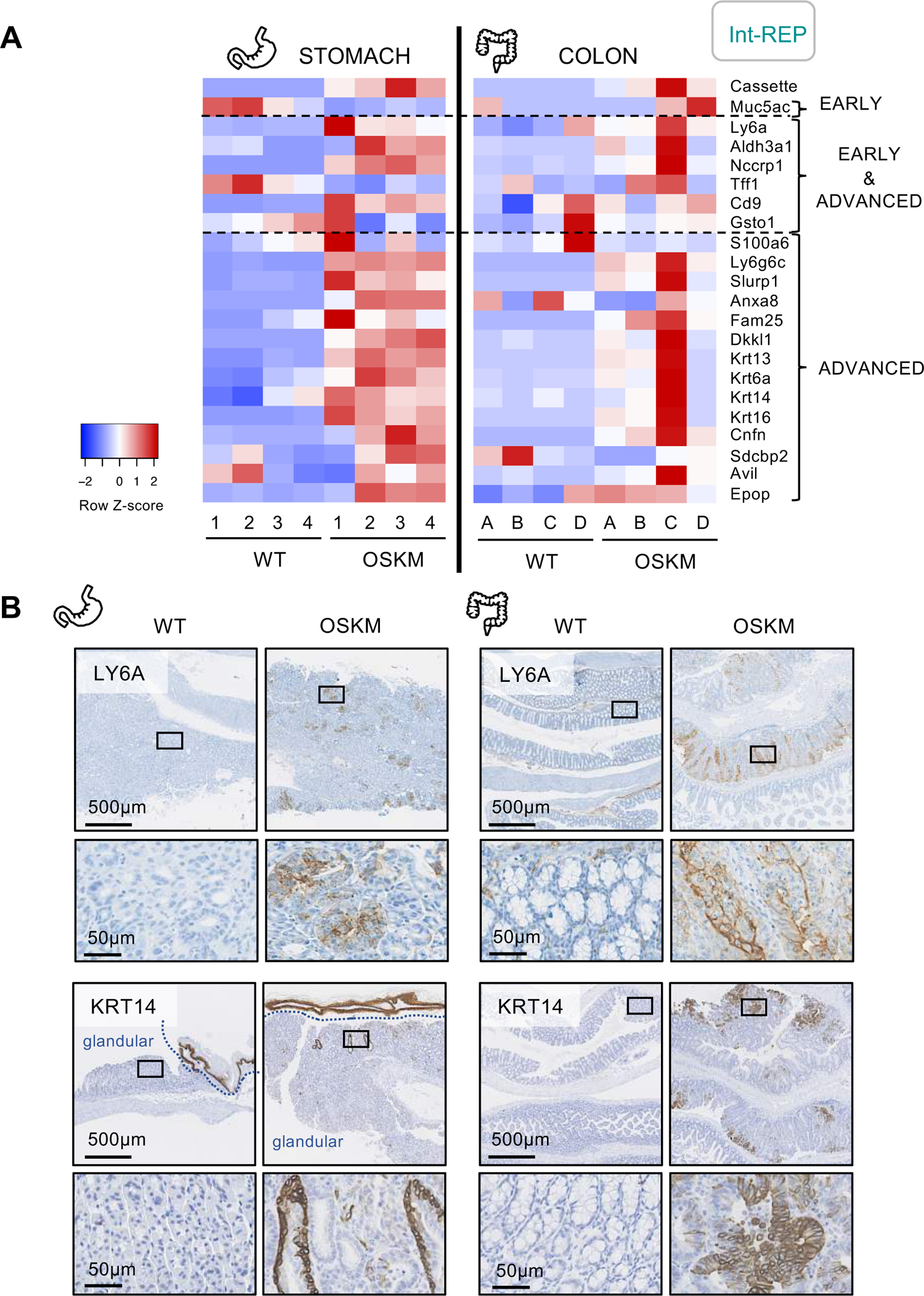
Markers of “intermediate reprogramming” are shared among several tissues. (A) Heatmap representation of mRNA expression by qRT-PCR of the identified markers of the “intermediate-reprogramming” state in stomach and colon samples of WT (n=4) and OSKM-induced (n=4) mice. Samples do not correspond to the same group of mice. Blue represents low expression and red high expression. (B) Immunohistochemistry of LY6A and KRT14 in intestine and stomach samples of WT and OSKM-treated mice. Scale bars, 500μm and 50 μm, as indicated on each image.

## Discussion

In this work, we report the first roadmap of *in vivo* partial reprogramming. Single cell RNA sequencing of reprogrammed pancreas suggests that pancreatic cell types are not equally affected by OSKM activation, with acinar cells being the most susceptible to reprogramming. This probably reflects the known plasticity of acinar cells, which can rapidly dedifferentiate upon inflammatory or oncogenic damages (van Roey et al., 2021). Also, expression of OSKM in acinar cells results in a general shut down of enhancers, and dedifferentiation (Shibata et al., 2018). In our analysis, we included a model of acute pancreatitis induced by intraperitoneal injections of cerulein for 2 consecutive days. We chose this control group for comparing OSKM-induced dysplasia with another type of pancreatic damage. It is well-established that acute pancreatitis caused by cerulein leads to the dedifferentiation of acinar cells in a process known as acinar-to-ductal metaplasia or ADM. Our scRNAseq analysis indicates that OSKM-reprogrammed acinar cells undergo a type of dedifferentiation that is distinct from cerulein-induced ADM. Regarding the rest of the pancreatic cell types, endocrine cells appear to be resistant to OSKM reprogramming, while ductal cells acquire a reactive molecular profile similar to ductal cells in cerulein-treated pancreas. It is possible that each cell type requires different periods of OSKM activation to surrender their somatic identity, probably reflecting the potency of epigenetic barriers (Arabacı et al., 2021).

Focusing on acinar cells, we found that a fraction of acinar cells still retain their somatic identity in the context of OSKM-reprogrammed tissue, while simultaneously acquiring a distinct molecular profile that we have called “pre-reprogramming” (pre-REP) and “non-reprogramming”, the latter one divided in two clusters non-REP and non-REP/AP-1. The pre-REP cluster, according to the trajectory analysis, is a hub that originates three different fates, namely, non-REP, non-REP/AP-1 and int-REP, being the latter the one giving rise to cells with pluripotency features. This situation is reminiscent of *in vitro* reprogramming in the sense that a substantial fraction of cells resist dedifferentiation. In contrast to normal acinar cells in non-reprogrammable mice, the three clusters that retain acinar identity during OSKM expression (pre-REP, non-REP and non-REP/AP-1) express high levels of the *Reg3* family, which are generally upregulated upon injury or stress in the pancreas (Chen et al., 2019). Regarding the non-REP/AP-1 cluster, it is worth mentioning that the AP-1 family of transcription factors plays a critical role during the early stages of *in vitro* reprogramming and successful progression towards pluripotency requires their subsequent downregulation (Chronis et al., 2017). Indeed, consistent with our observations, constitutive high expression of AP-1 factors, such as FRA1 or JUN, blocks progression of reprogramming (Chronis et al., 2017). In the case of the non-REP cluster, we identified two markers that were exclusive of this cluster, *Lrg1* and *Lcn2*, being absent in the pre-REP and the non-REP/AP-1 clusters. LRG1 has been reported as a biomarker for several malignancies, including pancreatic cancer (Fukamachi et al., 2019; Xie et al., 2019), LCN2 is a biomarker of tissue injury and metabolic disorders (Jaberi et al., 2021).

Senescent cells have been described to be generated during *in vivo* reprogramming and to play a key role in this process by providing secreted factors that promote reprogramming, such as IL6 (Mosteiro et al., 2016). However, we were not able to detect senescent cells (*p16/Cdkn2a*+ cells) in our scRNAseq analysis. We speculate that senescent cells may be under-represented due to their large size that can interfere with the process of cell isolation for 10x Genomics sequencing.

Regarding the fraction of acinar cells that succumb to OSKM-induced dedifferentiation, it consists of a heterogeneous population of cells that we have called “intermediate-reprogramming” (int-REP), which are the main focus of this study. This heterogeneous cluster includes very few cells expressing *Oct4*, suggesting that it contains cells on their way to pluripotency. Of note, our analysis has been done after 7 days of exposure to doxycycline, while full reprogramming usually requires a longer time frame in this animal model on the order of weeks (Abad et al., 2013). Based on the trajectory of cell identity change, we were able to distinguish early and advanced markers of intermediate reprograming, as well as markers that were present at both stages, early and advanced. Among the early markers of intermediate reprogramming we found *Muc5ac*, a mucin-encoding gene absent in normal pancreas but characteristic of the gastrointestinal epithelium. One of the most prominent markers that appears to be present in early and advanced intermediate reprogramming is *Ly6a* (also known as *Sca1*), a member of LY6 family associated with stem/progenitor cells in the hematopoietic system and in solid tissues such as the pancreas (Dzierzak and Bigas, 2018; Holmes and Stanford, 2007; Leinenkugel et al., 2022). It is remarkable that the large majority of dysplastic tissue is LY6A+ and indeed there is an excellent positive correlation between dysplasia and Ly6A-positivity. Until now, we could only estimate reprogramming efficiency by evaluating histological alterations. Here, we identify LY6A as an optimal marker for visualizing reprogrammed areas *in vivo*. Of note, *Ly6a* has been previously reported to be rapidly and transiently upregulated during *in vitro* reprogramming of mouse embryonic fibroblasts (MEFs) as well (Schwarz et al., 2018). Among the markers of advanced intermediate reprogramming, we found the upregulation of several keratins, including those characteristic of stratified epithelia, such as *Krt14*, a keratin that is not normally expressed in pancreas, except in the context of rare squamous pancreatic cancers (Bailey et al., 2016; Real et al., 1993).

We noted that the early marker of intermediate reprogramming, MUC5AC, corresponds to a protein that is normally absent in the pancreas, but it is characteristic of other tissues of endodermal origin, such as lung and stomach. On the other hand, the advanced intermediate reprogramming cells express keratins characteristic of epithelia of ectodermal origin, such as KRT14. Although highly speculative at present, this could suggest a trajectory of reverse developmental progression, from endoderm-like features to ectoderm-like features.

Regarding cancer, it has been reported that OSKM expression in the pancreas accelerates oncogenic KRAS-driven neoplasia (Shibata et al., 2018). In this regard, it is worth mentioning that some of the markers of intermediate reprogramming have been connected with pancreatic cancer. In particular, MUC5AC is abundantly expressed in the earliest neoplastic lesions of the pancreas, known as pancreatic intraductal neoplasia (PanIN) (Ganguly et al., 2021). Moreover, CD9 has been reported to be expressed and functionally relevant for pancreatic cancer stem cells (Wang et al., 2019). Finally, a fraction of advanced pancreatic ductal adenocarcinomas expresses KRT14 (Bailey et al., 2016; Real et al., 1993) and this has been mechanistically connected with the loss of GATA6, a key transcription factor for acinar identity (Martinelli et al., 2017).

It is now well-documented that *in vitro*, partially reprogrammed cells are heterogeneous and can acquire a variety of fates depending on the clues from the extracellular medium (see Introduction). In this regard, we wondered if the markers that we have identified in pancreas could be extrapolated to other tissues. Interestingly, most markers of pancreas intermediate reprogramming were upregulated in other tissues undergoing reprogramming, such as colon and stomach, as well as in reprogrammed fibroblasts (MEFs) *in vitro*. We also visualized partially reprogrammed colon and stomach using immunohistochemistry for LY6A and KRT14. This demonstrates the conservation of those markers among different tissue contexts and even among *in vitro* culture conditions. Overall, this work illustrates intermediates states of OSKM reprogramming *in vivo* and provides the first molecular roadmap of this process. Dissecting the states of *in vivo* reprogramming could be essential for future applications in tissue regeneration and cellular rejuvenation.

## Acknowledgements

We are grateful to Maria Isabel Muñoz for her assistance with the animal protocols and to Natalia del Pozo for valuable contributions. We thank the IRB core Facilities for their support, the Barcelona Science Park Animal Facility for animal maintenance, and the Spanish National Center of Genomic Analysis (CNAG) for the scRNA sequencing. D.C. was recipient of a fellowship from “la Caixa” Foundation. E.M was funded by a Future Fellowship from the IRB. M.K is supported by a fellowship from the Spanish Association Against Cancer (AECC). Work in the laboratory of F.X.R. was funded by grants from the Spanish Ministry of Science co-funded by the ERDF-EU (SAF2015-70553-R and RTI2018-101071-B-I00). Work in the laboratory of M.S. was funded by the IRB, and by grants from Spanish Ministry of Science co-funded by the ERDF-EU (SAF2017-82613-R), European Research Council (ERC-2014-AdG/669622), Secretaria d’Universitats i Recerca del Departament d’Empresa i Coneixement of Catalonia (Grup de Recerca consolidat 2017 SGR 282), “la Caixa” Foundation and the Milky Way Research Foundation. CNIO and IRB are supported by the Spanish Ministry of Science as Centres of Excellence “Severo Ochoa”.

## Authors Contributions

D.C. designed most experiments, performed animal experimentation, collected samples, performed RNA analysis, prepared the figures, and co-wrote the manuscript. J.M.d.V. performed all the bioinformatic analysis related to the single-cell RNA sequencing. E.M. contributed to the animal experimentation and collection of samples. C.L. performed the RNA-FISH. M.K. contributed to the animal experimentation and collection of samples. M.A. developed all the histological staining. N.P. supervised and interpreted the histological analyses. F.X.R. supervised the analysis of single-cell RNA sequencing analysis and provided his pancreas expertise throughout the study. M.S. designed and supervised the study, and co-wrote the manuscript. All authors discussed the results and commented on the manuscript.

## STAR Methods

### Key resources table

| REAGENT or RESOURCE | SOURCE | IDENTIFIER |
| --- | --- | --- |
| **Antibodies** |  |  |
| Mouse monoclonal anti-REG3A/G | Santa cruz biotechnology | CAT# 377038 |
| Rat monoclonal anti-LY6A | Abcam | CAT# 51317 |
| Rabbit polyclonal anti-CD9 | Abcam | CAT# 223052 |
| Rabbit polyclonal anti-KRT14 | Biolegend | CAT# 905301 |
| BrightVision poly HRP-Anti-Rabbit IgG | ImmunoLogic | CAT# DPVR-110HRP |
| Goat Anti-Mouse HRP | Dako - Agilent | CAT# P0447 |
| OmniMap anti-Rat HRP | Roche | CAT# 760-4457 |
| **Commercial assays** |  |  |
| Chromium Single Cell 3' GEM, Library & Gel Bead Kit v3 | 10X Genomics | PN-1000075 |
| BUNiScript™ cDNA Synthesis Kit for RT-qPCR | Bio-Rad | CAT# 170-8891 |
| iScript™ Advanced cDNA Synthesis Kit | Bio-Rad | CAT# 172-5038 |
| **Chemicals, peptides, and recombinant proteins** |  |  |
| Doxycycline hyclate BioChemica | PanReac | CAT# A2951 |
| Sucrose | Molecula | CAT# 90005036 |
| Cerulein | Bachem | CAT# 17650-98-5 |
| Dulbecco's modified Eagle's medium (DMEM) | Life Technologies | CAT# 31966047 |
| Fetal Bovine Serum (FBS) | Life Technologies | CAT# 10270106 |
| Penicillin-streptomycin | Gibco | CAT# 11528876 |
| KnockOut Serum Replacement (KSR) | Life Technologies | CAT# 10828028 |
| ESGRO Recombinant Mouse LIF protein | Merck Chemicals | CAT# ESG1107 |
| Non-essential Amino Acids solution (NEAA) | Life Technologies | CAT# 11140035 |
| β-mercaptoethanol | Life Technologies | CAT# 31350010 |
| Doxycycline | PanReac | CAT# A2951 |
| Collagenase P | Sigma | CAT# 1121386500 |
| Dispase II | Life Technologies | CAT# 17105041 |
| Soybean Trypsin Inhibitor | Life Technologies | CAT# 17075-029 |
| DNase I | Sigma | CAT# D4513 |
| HBSS with Ca<sup>2+</sup>/Mg<sup>2+</sup> | Life Technologies | CAT# 14025050 |
| Trypsin-EDTA | Life Technologies | CAT# 25300062 |
| Red blood cell lysis buffer | Biolegend | CAT# 42030 |
| Trizol | Life Technologies | CAT# 15596018 |
| Go-Taq qPCR Master Mix | Promega | CAT# A6002 |
| 10% neutral buffered formalin | Sigma-Aldrich | CAT# HT501128-4L |
| Formamide | Merck | CAT# 344205 |
| Proteinase K | Sigma-Aldrich | CAT# 3115879001 |

| **Software and algorithms** |  |  |
| --- | --- | --- |
| Seurat R package |  | v. 4.1.0 |
| SoupX |  | v. 1.5.2 |
| Slingshot |  | v. 1.8.0 |
| NDP.view 2 |  | v. 2 |
| Picky 2.0 |  | v. 2.0 |
| ImageJ |  | v. 1.53K |
| QuPath software |  | v. 0.1.2 |
| GraphPad Prism |  | v. 9.0.1 |

### Resource availability

#### Lead contact

Further information and requests for resources and reagents should be directed to and will be fulfilled by the lead contact, Manuel Serrano.

### Materials availability

This study did not generate new unique reagents.

### Experimental models and subject details

#### Ethics Statement

Animal experimentation was performed between two different institutes: at the Spanish National Cancer Research Centre CNIO in Madrid and at the Institute of Research in Biomedicine IRB in Barcelona, according to protocols approved by the CNIO-ISCIII Ethical Committee for Research and Animal Welfare (CEIyBA) in Madrid, and by the Animal Care and Use Ethical Committee of animal experimentation of Barcelona Science Park (CEEAPCB) and the Catalan Government in Barcelona.

#### Mouse model

We used the reprogrammable mice known as i4F-B which carries a ubiquitous doxycycline-inducible OSKM transgene, abbreviated as i4F, and inserted into the Pparg gene (Abad et al., 2013). Mice of both sexes were used for the experiments.

## Method details

### Animal treatments

To activate the four Yamanaka factors, 1 mg/ml of Doxycycline hyclate BioChemica (PanReac, A2951) was administered in the drinking water supplemented with 7.5% sucrose for a period of 7 days and mice were sacrificed directly after.

Acute pancreatitis was induced as previously described (Carrière et al., 2011). In brief, adult wild-type (WT) mice were intraperitoneally injected with cerulein (Bachem), an analog of the pancreatic secretagogue cholecystokinin for mice, dissolved in saline solution (100ug/kg body weight) hourly for 7 hours on two consecutive days. Mice were sacrificed the day after the last injection.

### Cell culture

Primary mouse Embryonic Fibroblasts (MEFs) were obtained from i4F embryos at embryonic stage E13.5 and cultured in Dulbecco’s modified Eagle’s medium (DMEM) (Gibco) supplemented with 10% of heat-inactivated Fetal Bovine Serum (FBS) (Life Technologies) and penicillin-streptomycin (Gibco). For *in vitro* reprogramming, 3 x 10^6^ i4F MEFs were seeded in 6 well-plates (Corning) and cultured with “IPSC medium”, composed by high glucose DMEM (Gibco) supplemented with 15% of KnockOut Serum Replacement (KSR) (Life Technologies), LIF (1000 U/ml) (Merck Chemicals, Germany), non-essential amino acids, penicillin-streptomycin (Gibco) and 100 μM β-mercaptoethanol (Life Technologies, 31350010). Cells were treated with 1 μg/ml of doxycycline (PanReac, A2951) to induce the expression of OSKM cassette. Medium was changed every 48 hours until IPSC colonies emerged approximately at day 11. IPSCs were cultured on irradiated (20 rads) feeder MEFs on gelatin-coated plates with “IPSC medium”.

### Tissue processing and single-cell RNA sequencing

Mouse primary pancreatic cells were obtained by digesting the whole pancreas with 1 mg/mL Collagenase P (Sigma, 11213865001) supplemented with 2 U/mL Dispase II (Life Technologies, 17105041), 0.1 mg/mL Soybean Trypsin Inhibitor (Life Technologies, 17075-029) and 0.1 mg/mL DNase I (Sigma, D4513) in HBSS with Ca^2+/^Mg^2+^ (Life Technologies, 14025050). Tissue was dissociated using the gentle MACS^TM^ Octo Dissociator (Miltenyi Biotech) at 37°C for 40 min, was further digested with 0.05% Trypsin-EDTA (Life Technologies, 25300062) for 5 min at 37°C and erythrocytes were removed by incubating the cell suspension with red blood cell lysis buffer (Biolegend, 420301, San Diego, US) for 5 min at RT. Single cell suspension of pancreatic cells was resuspended in FACs buffer (10mM EDTA, 2% FBS in Ca^2+/^Mg^2+^-free PBS) and DAPI negative cells were selected by cell sorting using FACSAria Fusion (BD Biosciences). Cell viability was above 70% for each sample. Sorted cells were loaded onto a 10xChromium Single Cell Controller chip B (10x Genomics) as described in the manufacturer’s protocol (Chromium Single Cell 3ʹ GEM, Library & Gel Bead Kit v3, PN-1000075). Generation of gel beads in emulsion (GEMs), barcoding, GEM-RT clean up, cDNA amplification, and library construction were performed following the manufacturer’s recommendations. Libraries were loaded at a concentration of 1.8 pM and sequenced in an asymmetrical pair-end format in a NextSeq500 instrument (Illumina).

### scRNAseq data analysis

Reads were demultiplexed and aligned to mm10 genome using Cell Ranger software. Filtered and unfiltered resultant matrices were used as input for SoupX library (v.1.5.2) (ref.below) to estimate ambient RNA contamination. Top contaminant genes were removed from the count matrices (Ctrb1, Prss2, Cela2a, Cela3b, Cela1, Pnlip, 2210010C04Rik, Try5, Try4, Clps, Ctrl, Cpa1,Cpb1, Sycn, Rnase1, Zg16, Reg1, Ins2, Pnliprp1, Cel, Cpa2, Ins1, Klk1, Tff2, Gcg) (Young and Behjati, 2020). Cell Ranger matrix data (barcodes, features, and count matrix) were loaded onto the Seurat R package (version 4.1.0.). Cells expressing >5% of mitochondrial genes were excluded from downstream analysis, considered as low-quality cells. Normalization and variance stabilization of filtered cells was performed using regularized negative binomial regression (SCTransform) (Hafemeister and Satija, 2019). Linear dimensional reduction was performed on the normalized data using principal component analysis (PCA). Elbow plots and JackStraw permutations test were applied to determine the dimensionality of each dataset as significant principal components. The identification of biologically relevant communities was performed by means of a graph-based clustering in Seurat through the construction of a KNN graph and posterior cell clusterization using the Louvain algorithm (resolution 0.2). Cell clusters were visualized using uniform manifold approximation and projection (UMAP) (Becht et al., 2018) plots with previously selected significant components as an input. Cluster gene markers were detected with Seurat R package FindAllmarkers function using a Wilcoxon rank sum test between each cluster and the rest of cells in the dataset and p value adjustment was performed using Bonferroni correction based on the total number of genes in the dataset. Pseudotime trajectory analysis was performed on the Seurat object using Slingshot package (v 1.8.0). Briefly, the global lineage structure was identified by constructing the minimum spanning tree on the clusters with the getLineages function and then smooth lineages and pseudotime variables were inferred by fitting simultaneous principal curves with the getCurves function.

### RNA isolation and Analysis of mRNA levels

Total RNA was extracted from cells and tissues such as colon and stomach with Trizol (Invitrogen), following provider’s recommendations, and 1 μg of RNA was reverse transcribed into cDNA using BUNiScriptTM cDNA Synthesis Kit for RT-qPCR (Bio-Rad, 170-8891). For pancreas samples, total RNA was isolated using guanidine thiocyanate, followed by acid phenol-chloroform extraction, and up to 5 μg of total RNA was reverse transcribed into cDNA using iScript^TM^ Advanced cDNA Synthesis Kit for RT-qPCR (Bio-Rad, 172-5038). Quantitative real time-PCR was performed using Go-Taq qPCR Master Mix (Promega) in a QuantStudio 6 Flex thermocycler (Applied Biosystem). For input normalization, we used the housekeeping gene *Gapdh* and *β-Actin*. The primers used are listed in the following table (**Table 1**):

**Table 1:**
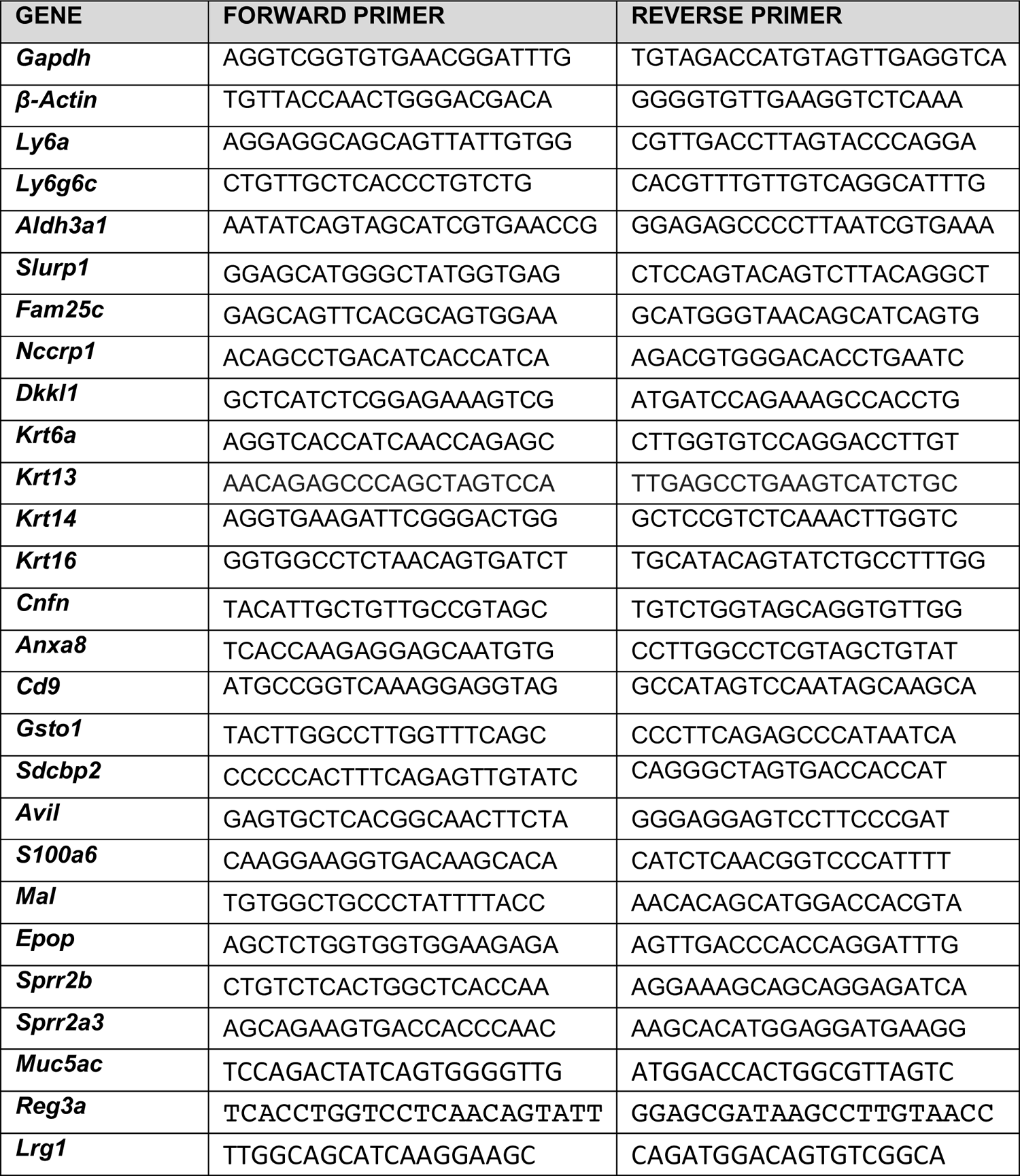
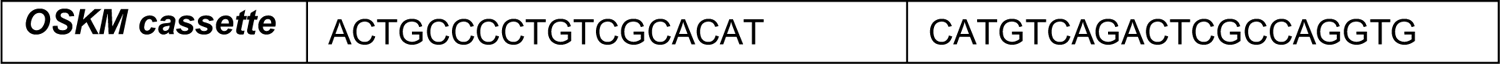
Mouse Primers for RT-qPCR

### Histology and Immunohistochemical study

Tissues were fixed overnight in 10% neutral buffered formalin (HT501128-4L, Sigma-Aldrich), embedded in paraffin blocks, sectioned at a thickness of 3 μm, mounted in Superfrost^®^plus slides and dried. Serial tissue sections were deparaffinized in xylene and rehydrated through a series of graded ethanol until water, and then stained with hematoxylin and eosin (HE) for the visualization of the tissue architecture, as well as with different antibodies (see **Key resources table**). In most of the cases, immunohistochemistry was performed by an automated immunostaining platform (Ventana discovery XT, Roche). If not optimized for automatization, the staining was performed manually following standard protocols. As a general guideline, antigen retrieval was mainly performed with high pH buffer (CC1m, Roche), endogenous peroxidase was blocked and slides were then incubated with the appropriate primary antibodies as detailed (see **Key resources table**). After the primary antibody, slides were incubated with the corresponding secondary antibodies and visualization systems (OmniMap, Ventana, Roche) conjugated with horseradish peroxidase (Chromomap, Ventana, Roche). Immunohistochemical reaction was developed using 3,3-diaminobenzidine tetrahydrochloride (DAB) as a chromogen and nuclei were counterstained with hematoxylin. Finally, the slides were dehydrated, cleared and mounted with a permanent mounting medium for microscopic evaluation. Whole digital slides were acquired with a NanoZoomer S60 Digital slide scanner, and images captured with the NDP.view2 Viewing software.

### Image analysis

Brightfield images of immunohistochemistry were quantified in a blinded way using QuPath software 0.1.2 with standard DAB detection methods. OSKM-induced dysplasia was evaluated and quantified by histopathological assessment of those areas in the pancreas.

### RNA fluorescence *in situ* hybridization (RNA-FISH)

In situ 4-colour RNA-FISH was performed using primer-padlock 2-oligo hybridization of the RNA targets (Table S4), essentially as described (Wang et al., 2018), except that hybridization was performed in 30% formamide buffer (Merck). Briefly, ssDNA oligo probes were designed against the coding regions of target RNA transcripts (with the exception of the OSKM cassette, where the non-coding regions were deliberately selected), using Picky2.0 to identify loci without RNA secondary structure or repetitive sequence, followed by blast searching to confirm target specificity. Probe hybridization was performed overnight at 40°C with rocking in a humid chamber. Probes then underwent rolling circle amplification, polyacrylamide gel mounting and proteinase K (Sigma-Aldrich) sample clarification before imaging, images were acquired in a Zeiss LSM880 microscope, with processing in ImageJ using background subtraction (50 pixels) and a 2-pixel median-filter.

### Statistical analysis

The data were analyzed using GraphPad Prism v.9.0.1 software. Linear regression was using to correlate the expression of markers (LY6A and KRTA14) with the extent of the dysplasia in the pancreas.

### Data and code availability

Generated scRNA-seq datasets have been deposited in GEO under accession number GSE188819.

---

**Figure S1, Related to Figure 1.**
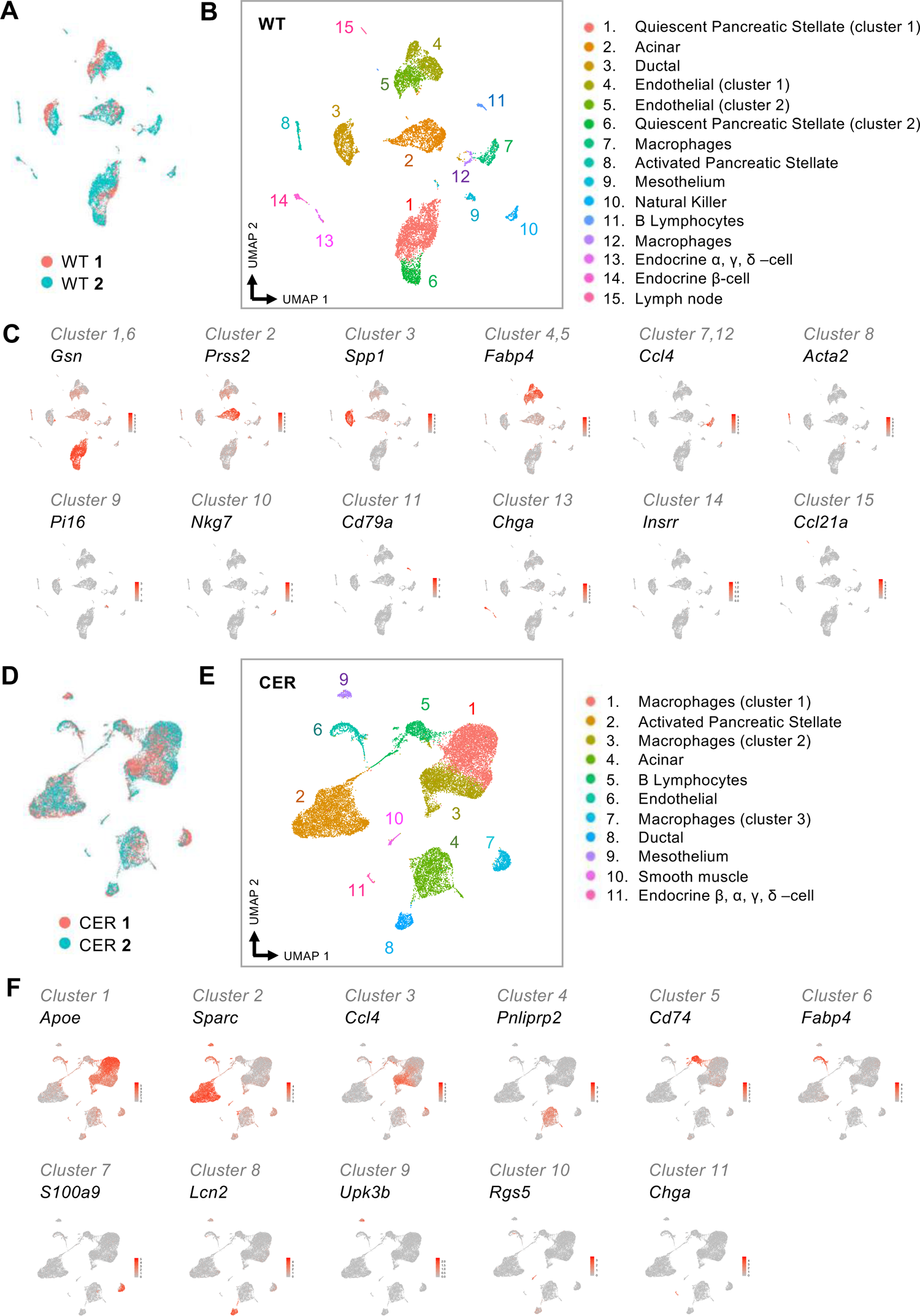
Single cell RNA sequencing of the whole pancreas of WT and CER-treated mice. (A) UMAP (Uniform Manifold Approximation and Projection) visualization of two wild-type pancreata (WT 1 and WT 2) after merging them. (B) UMAP visualization of the 15 populations emerging after clustering together with their annotations. (C) Representative marker used for the annotation of each cluster is depicted in a UMAP. (D) UMAP visualization of two cerulein (CER)-treated pancreata (CER 1 and CER 2) after merging them. (E) UMAP visualization of the 11 populations emerging after clustering together with their annotations. (F) Representative marker used for the annotation of each cluster is depicted in a UMAP.

**Figure S2, Related to Figure 1.**
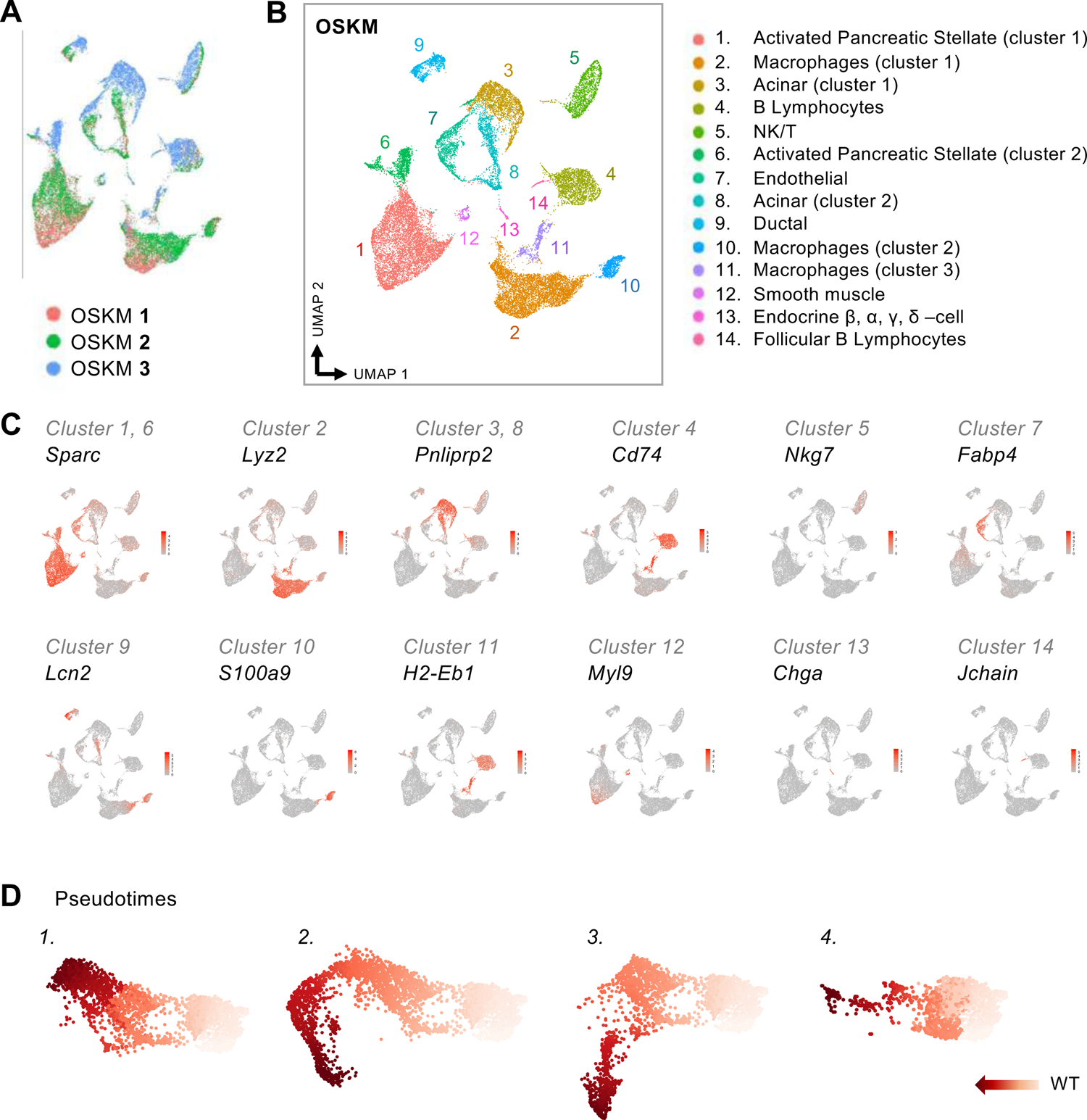
Single cell RNA sequencing of the whole pancreas coming from OSKM-induced mice. (A) UMAP visualization of three OSKM-induced pancreata (OSKM 1, OSKM 2 AND OSKM 3) after merging them. (B) UMAP visualization of the 14 populations emerging after clustering together with their annotations. (C) Representative marker for each cluster is depicted in a UMAP representation. (D) WT acinar cells were pre-selected as the starting point of the trajectory analysis. Pseudotimes 1 to 4 illustrate each cell lineage separately with light color to mark the start and the color gradient the transcriptional progression of each lineage to its terminal state.

**Figure S3, Related to Figure 2.**
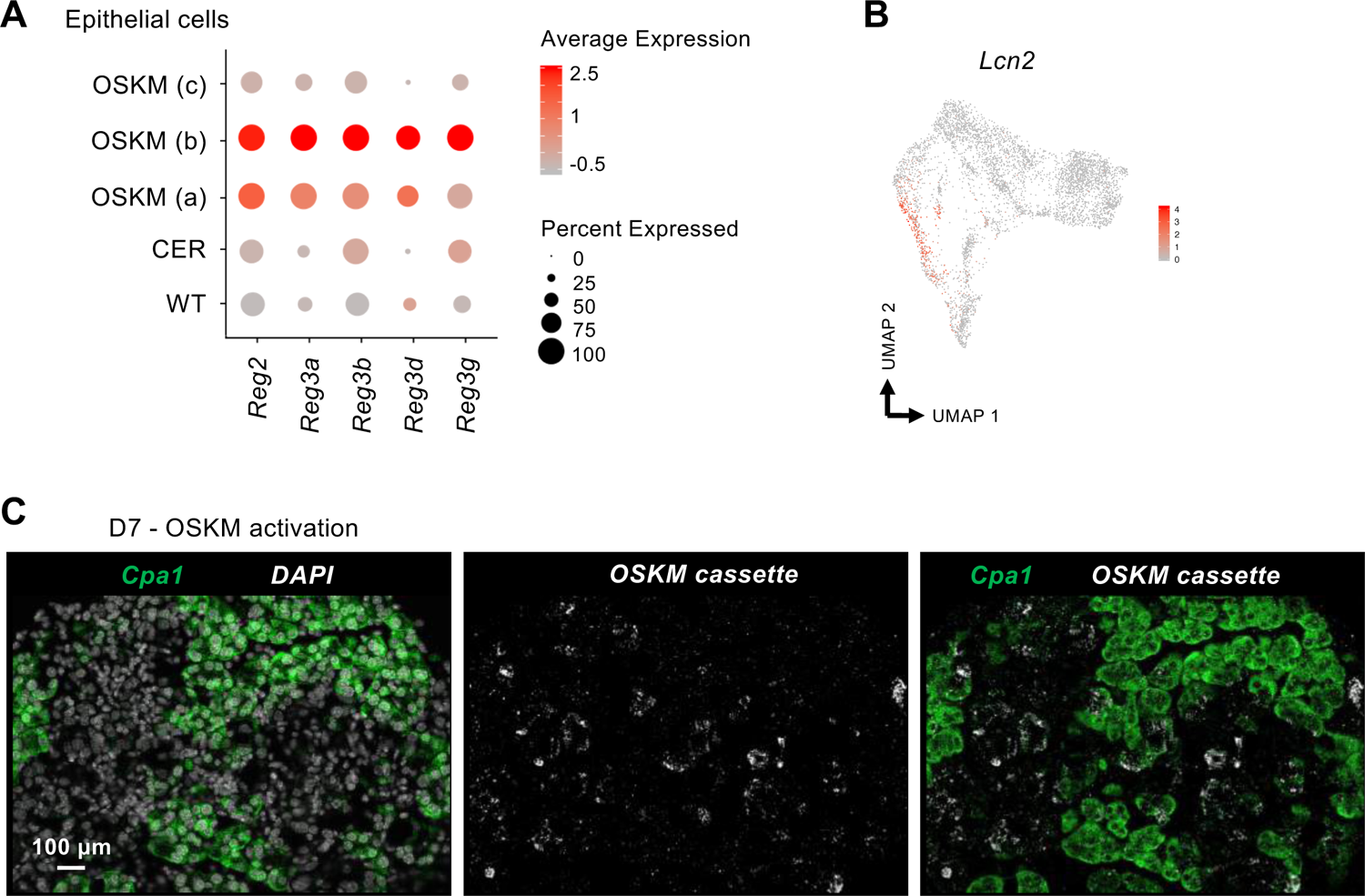
*In vivo* identification of markers characterizing “pre-reprogramming” and “non-reprogramming” OSKM fates. (A) Expression of various members of *Reg* family in epithelial cells coming from all three conditions. (B) UMAP visualization of *Lcn2* and *Itih4* expression in acinar cells coming from WT and OSKM-induced mice. (C) Images of RNA-fluorescence *in situ* hybridization (RNA-FISH) of partially reprogrammed pancreas (day 7 of OSKM activation) using probes against *Cpa1* and *OSKM cassette* together with DAPI. Scale bar, 100 μm.

**Figure S4, Related to Figure 3.**
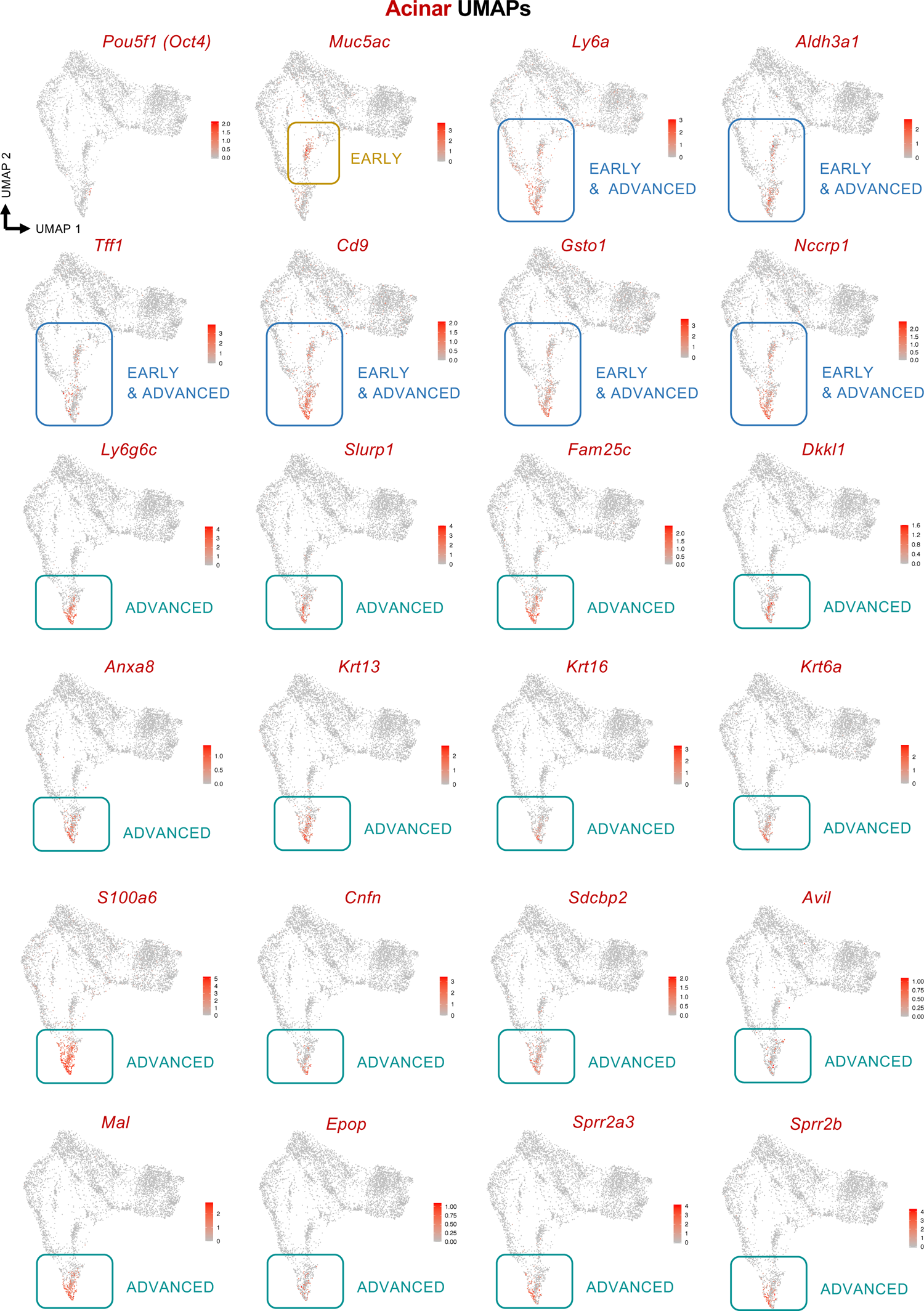
*In vivo* identification of “intermediate-reprogramming” fate. UMAP visualization of all selected genes that characterize cells undergoing OSKM-reprogramming: the pluripotent marker *Pou5f1* (*Oct4*), “early” markers of intermediate-reprogramming (*Muc5ac*), “early & advanced” markers (*Ly6a, Aldh3a1, Tff1, Cd9, Gsto1, Nccpr1*) and “advanced” markers (*Ly6g6c, Slurp1, Fam25c, Dkkl1, Anxa8, Krt13, Krt16, Krt6a, S100a6, Cnfn, Sdcbp2, Avil, Mal, Epop, Sprr2a3, Sprr2b*)

**Figure S5, Related to Figure 4.**
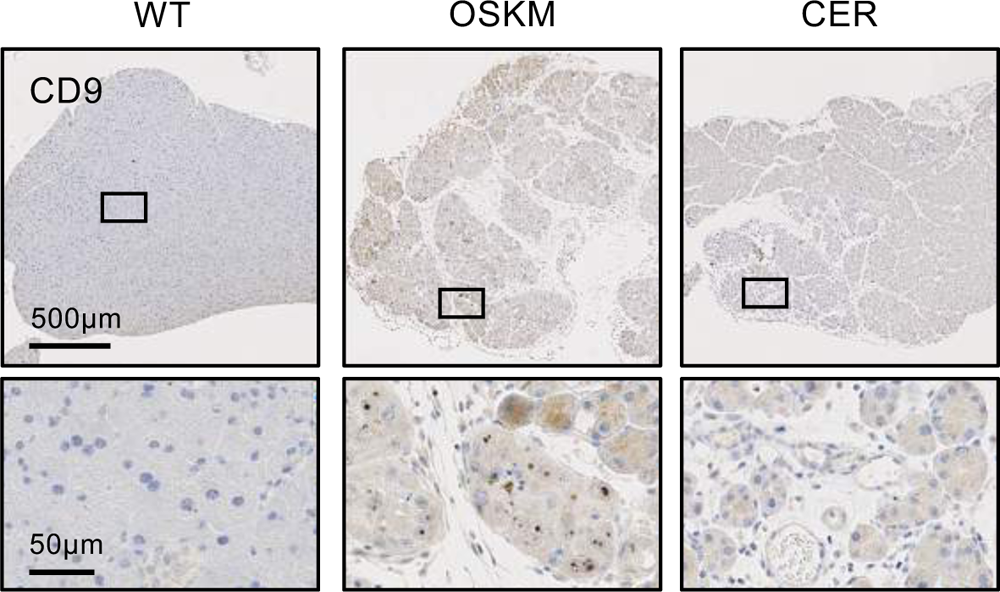
*In vivo* identification of “intermediate-reprogramming” fate. Immunohistochemistry of CD9 in the pancreas of WT, CER-treated and OSKM-induced mice.

**Figure S6, Related to Figure 7.**
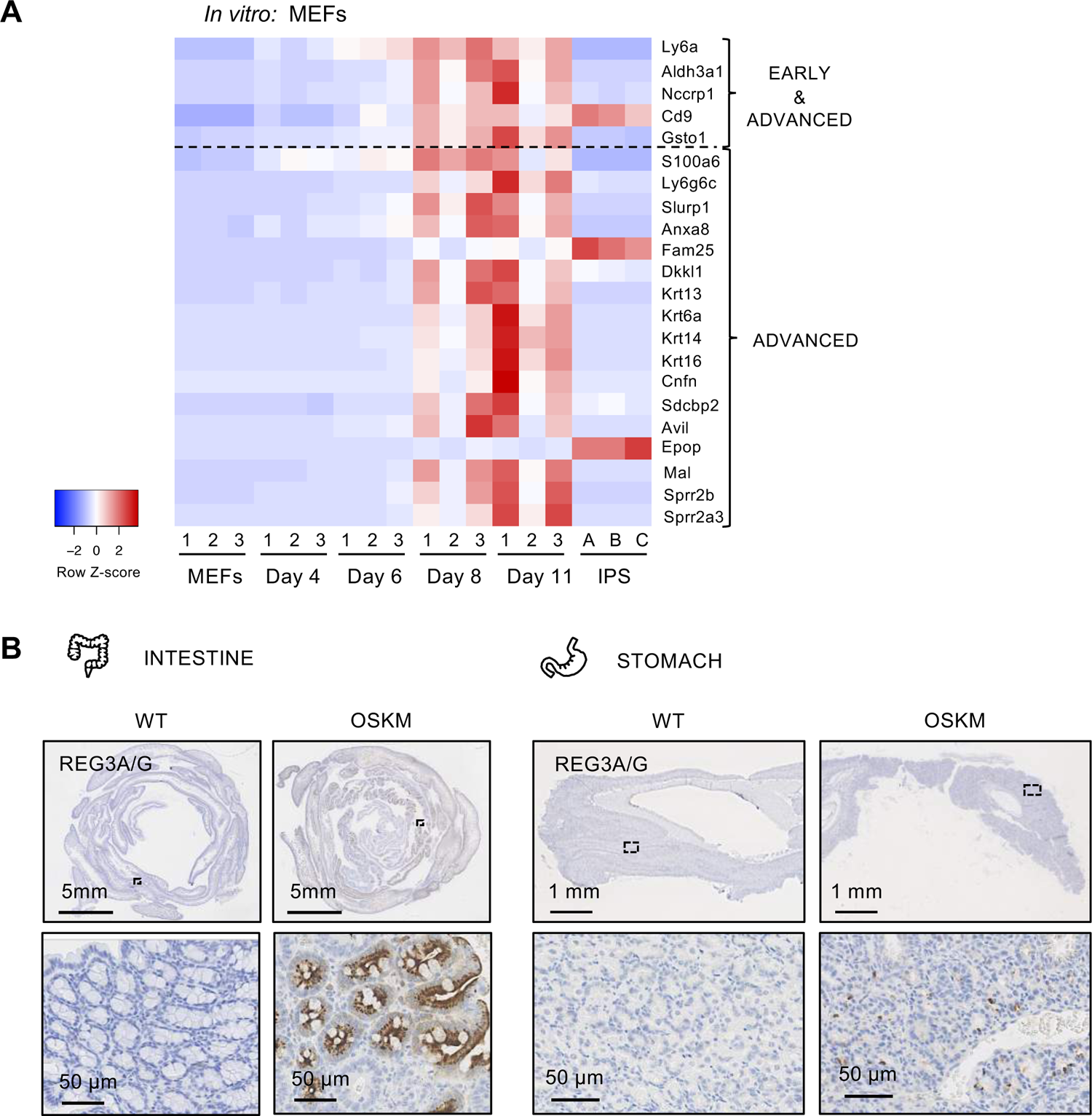
Markers of *in vivo* reprogramming are also present during *in vitro* reprogramming of MEFs and in other tissues. (A) Heatmap representation of mRNA expression of the identified markers for “intermediate-reprogramming” state in mouse embryonic fibroblasts (MEFs) undergoing reprogramming *in vitro,* as well as induced pluripotent stem cells (IPS). Blue represents low expression and red high expression. (B) Immunohistochemistry of REG3A/G in intestine and stomach samples coming from WT and OSKM-treated mice. Scale bars, 5mm, 1mm and 50μm, as indicated on each image.

